# The mineralization of osteonal cement line depends on where the osteon is formed

**DOI:** 10.1101/2024.10.06.616843

**Authors:** A. Cantamessa, S. Blouin, M. Rummler, A. Berzlanovich, R. Weinkamer, M.A. Hartmann, D. Ruffoni

**Affiliations:** Mechanics of Biological and Bioinspired Materials Laboratory, Department of Aerospace and Mechanical Engineering, University of Liège, Liège, Belgium; Ludwig Boltzmann Institute of Osteology at Hanusch Hospital of OEGK and AUVA Trauma Centre Meidling, 1^st^ Medical Department Hanusch Hospital, Vienna, Austria; Department of Biomaterials, Max Planck Institute of Colloids and Interfaces, Potsdam, Germany; Center of Forensics Medicine, Vienna, Austria

**Keywords:** cement line, mineralization, osteon, human bone, quantitative backscattered electron imaging, mineralization kinetics, mineral content, mineral recycling, tissue age

## Abstract

The cement line (CL) is a thin layer separating secondary osteons from interstitial bone and other osteons. It is assumed to play a significant role in bone fracture resistance, owing to its ability to deflect or arrest microcracks. Despite the possible role for bone quality, the CL is still one of the least understood microstructural features of bones, with unknowns on CL composition, mineralization, and mechanical properties. This study, focusing on CL mineralization, aims to elucidate the interplay between the mineral content of the CL and of adjacent bone tissue. Using quantitative backscattered electron imaging, osteons with different degrees of mineralization coming from human femoral samples were analyzed. We implemented a spatially resolved analysis of the mineral content in layers along the CL, considering both regions inside the osteon (i.e., formed soon after CL deposition) and outside (i.e., already present at the time of CL deposition). We found that the mineral content of the CL correlates strongly with the mineral content outside of the osteon, but not inside. Assuming the mineral content of the osteon as a proxy of its age, we demonstrate that not only the osteon, but also the CL increases its mineral content with time. However, the rate of increase is lower in the CL. Importantly, the specific value of the high initial mineral content of the CL depends on the mineral content of the local surrounding, in which the osteon was formed. Our findings highlight the central role of the local degree of mineralization of the bone around the osteon for building the CL.

## Introduction

Cement lines (CLs) are thin interphases, typically 1-3 *µ*m in width, which are ubiquitous in bone and usually found at: i) the border of secondary osteons in cortical bone^(1)^, ii) around individual bone packets in trabecular bone^(2)^ and iii) between bone and mineralized cartilage at entheses^(3,4)^ and osteochondral junctions^(4)^. Despite their small thickness, CLs are assumed to contribute to bone fracture resistance by deflecting or arresting microcracks. In cortical bone, studies have shown that microcracks coming from the interstitial tissue, can - depending on their dimensions and paths - deviate when meeting the CL at the outer border of an osteon^(5–8)^. Such a mechanism does not only allow to dissipate energy and to reduce the force driving crack propagation^(9)^, but also to preserve the integrity of the osteon and of the corresponding blood vessels housed in the central Haversian canal. Indeed, microcracks are more likely observed in interstitial bone than inside osteons^(10)^. The ability of the CL to interact with damage depends on its basic mechanical properties such as stiffness, strength and toughness^(11)^. Although those properties are by far not well characterized, CL mineral content is one factor of fundamental importance for mechanical behavior. Yet, the composition of CLs remain controversial: both lower^(12)^ and higher^(13)^ mineral content of CLs compared to surrounding bone are reported. More recent data based on backscattered electron imaging, Raman spectroscopy and synchrotron radiation tomography indicate that CLs tend to have higher mineral content than the corresponding osteons^(10,14,15)^, and that this difference in mineral content decreases with tissue age^(15)^. At the same time, the dimensions of the osteon do not seem to influence the degree of mineralization of the CL^(15)^.

The mineral content of the osteon is not static but evolves in time according to specific mineralization kinetics: current understanding is that a fast primary mineralization, bringing the mineral content of newly deposited osteoid up to about 70% of the final value within a few days, is followed by a much slower secondary mineralization, leading to a gradual maturation of the mineral phase, which takes up to several years to complete^(16–19)^. Conversely, knowledge about how much mineral is incorporated during CL formation and to which extent the mineral content of the CL evolves with time is insufficient. Secondary osteons and CLs are a result of bone remodeling^(20)^, with CLs providing a proper “cleaned” surface for osteoid deposition^(1,21,22)^. CLs have higher content of osteopontin (a CL marker) than adjacent bone^(22)^. This protein can be produced by osteoclasts, osteoprogenitors cells as well as osteoblasts^(23)^. The examination of cutting cones in osteons with serial sectioning has provided clues that CLs are formed during or immediately after bone resorption by osteoclasts/osteoprogenitor cells depositing a matrix rich in osteopontin^(1)^. Much less osteopontin is seen in lamellar bone, which is deposited at the end of the reversal phase, once a critical density of osteoblasts has been reached^(1)^. In addition to osteopontin, a further element that can impact tissue mineralization is the transport of mineral ions to the mineralization site. There is evidence that locations of bone modeling (i.e., bone formation occurring on resting periosteal surfaces) and remodeling (i.e., bone formation preceded by resorption and occurring on eroded surfaces) present differences in the mineralization process, resulting in specific mineral characteristics such as crystal composition and dimensions^(24)^. This is likely due to different avenues to transport ionic precursors: usually bone modeling requires transport over large distances and, thus, ionic precursors should be stabilized by compositional modifications, whereas in bone remodeling this is not necessary as resorbed mineral may be locally recycled^(24)^. As the formation of CL and osteonal bone are somewhat dephased, both in space and time, differences in available mineral ions may be present, impacting the mineralization process.

The aim of this work is to provide a quantitative description of the mineral content of CLs taking into consideration the surrounding of the osteon and to infer from that data how this mineral content evolves in time. We explored potential relations between the mineral content of CLs and of adjacent bone tissue, considering regions located immediately outside the osteon, therefore close to the eroded surface and already present at the time of CL formation, as well as inside the osteon, i.e. deposited after the CL. We carried out a local analysis of the mineral content at submicrometer resolution using quantitative backscattered electron imaging (qBEI), considering several human osteons having different degrees of mineralization, which is assumed as an approximate descriptor of tissue age. By analyzing osteons of different ages, we aim to assess the mineralization kinetics of the CL.

## Materials and Methods

### Sample preparation

Human femurs from two male donors (40 and 81 years old) without known metabolic bone disease were obtained from the Department of Forensic Medicine of the Medical University of Vienna, in accordance with the ethical commission of the institution (EK no. 1757/2013). After extraction, the bones were kept frozen at -20 C until further preparation. Cross sections (perpendicular to the longitudinal axis the femurs) of about 1.5 cm in thickness were cut from the mid-diaphysis (Figs. 1A and B) using a water-cooled diamond saw (Buehler Isomet 1000, Germany). The sections were then prepared following a well-established procedure^(25,26)^, which involved dehydration in graded ethanol series, followed by immersion in ethanol/acetone baths, and embedding in poly-methyl methacrylate (PMMA). The transverse surfaces were ground with sandpapers having decreasing grit size (P1200, P2400) and polished using a diamond suspension down to 1 *µ*m particle size on a silk cloth under glycol irrigation (Logitech PM5, UK), to minimize the formation of microcracks^(27)^.

**Fig. 1:**
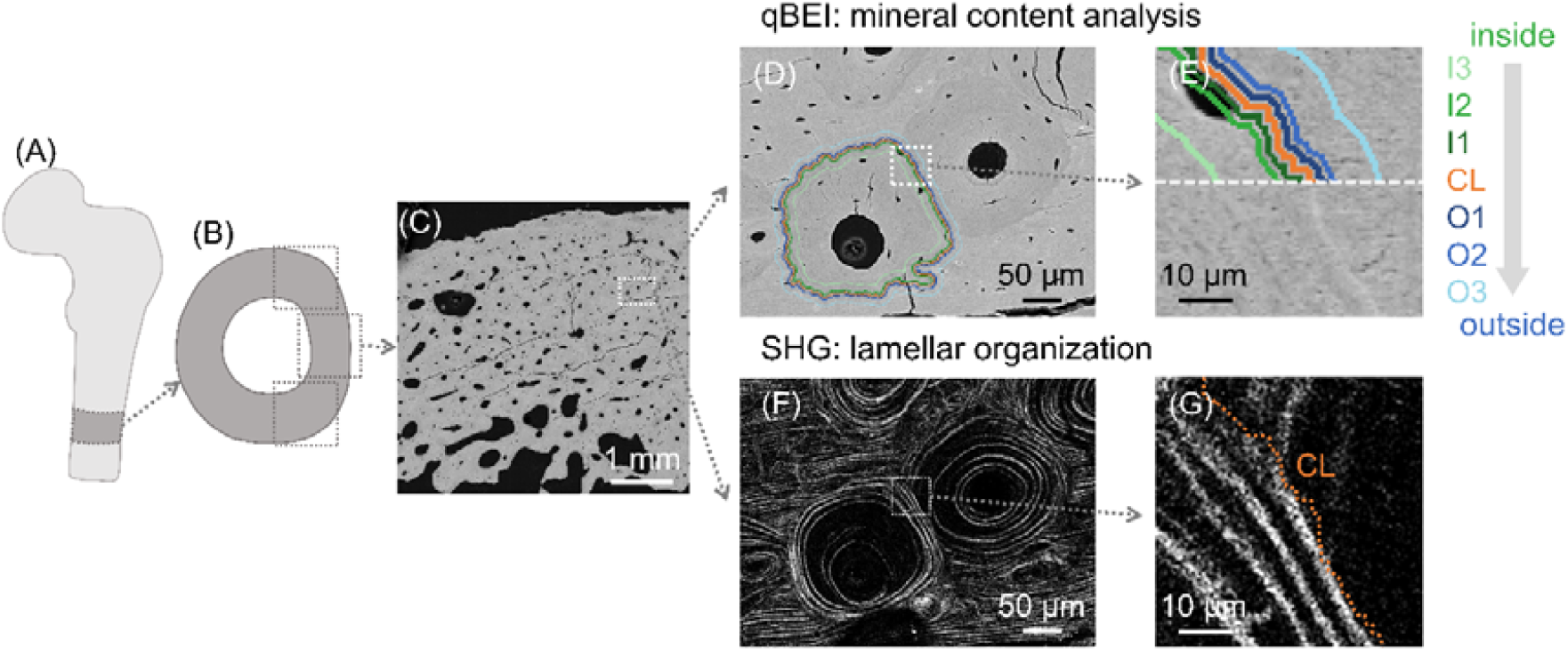
Analysis of human osteonal bone. (A) Schematic of a human femur and (B) transverse cross-section where (C) overview quantitative backscattered electron imaging (qBEI) scans were performed to select uninterrupted osteons. (D) High resolution qBEI map of a representative uninterrupted osteon neighboring older osteons (left and right) and interstitial bone (top and bottom). (E) Magnified view of a cement line (CL) and the different concentric layers used to analyze the mineral content around the CL. Layers are located both inside (I1, I2, I3) as well as outside (O1, O2, O3) the selected osteon with different distance to the CL. (F) Second harmonic generation (SHG) image of the same section analyzed with qBEI highlighting the lamellar organization. This information is used to ensure that the selected osteon is uninterrupted (i.e. the latest deposited in the neighborhood). (G) Close-up of the SHG image with the orange dashed line representing the location of the CL, detected by superimposing SHG and qBEI maps.

### Quantitative backscattered electron imaging (qBEI)

The local mineral content of cortical bone was quantified with qBEI, a well-established technique based on scanning electron microscopy (SEM), providing quantitative two-dimensional spatial maps of calcium content by measuring the intensity of the backscattered signal^(26,28)^. To obtain a conducting surface, a thin carbon layer was deposited on the polished samples by vacuum evaporation (Agar SEM carbon coater, Agar, Stansted, UK). The SEM, equipped with a zirconium-coated field emission cathode (Zeiss SEM SUPRA 40, Germany), was operated at 20 kV with a working distance of 10 mm, and a probe current of 280 – 320 pA. First, overview scans (Fig. 1C) with a larger field of view and a lower resolution (1.76 *µ*m/pixel) were performed to select the osteons of interest for subsequent higher resolution analysis. The selected osteons had to feature a central Haversian canal and a continuous border i.e., they should not be interrupted by more recent osteons. In the analysis, we included osteons with different size and mineral content, which is assumed as a proxy of tissue age. The chosen osteons (n = 35) and the surrounding bone were measured at higher magnification (pixel resolution 0.57 μm with a field of view of 1024 x 768 pixels). This resolution is high enough to resolve the CLs, which have a thickness in transverse sections of about 1-3 μm and low enough to allow for a quantitative measurement of the calcium content without additional contrast due to, e.g., orientation of bone lamellae. A calibration of the SEM backscattered signal with carbon and aluminum standards was performed as previously reported^(26,29)^, to establish a quantitative relation between the measured grey level and the local mineral content, expressed as calcium weight percent (Ca wt %).

### Second harmonic generation imaging

To ensure that the selected osteons were uninterrupted, which can be challenging when two adjacent osteons with similar mineral content slightly overlap, and to have qualitative information on the lamellar pattern of the osteons, second harmonic generation (SHG) imaging was performed on the same locations measured with qBEI (Figs. 1F and G). Samples were imaged using confocal laser scanning microscopy (Leica TCS SP8 DLS, Germany) with a 40x oil immersion objective (HC PL APO 40×/1.30 OIL). A pulsed infrared laser (Spectra-Physics, Milpitas) operated at a wavelength of 910 nm was used, and the backward direction signal was detected at 450–460 nm wavelength. Images were acquired with a nominal pixel size of 379 nm and a field of view of 1024 x 1024 pixels at a scan rate of 400 Hz. Several regions were analyzed and stitched together.

### Mineral content of the cement line and adjacent regions: layers analysis

Before evaluating the mineral content, a 2-pixel erosion was applied to minimize boundary effects^(16)^ using the software CTAn (v1.19.4.0, Skyscan). Background was removed by discarding all pixels with Ca-content below 5.2 wt %, corresponding to embedding resin and soft tissues. The mineral content of the CL and of neighboring bone was evaluated according to the following layer analysis, implemented in CTAn. First, a mask of each osteon (not including the CL) was generated by manually contouring the osteon border as delimited by the CL (Figs. S1A and B). This choice is motivated by the fact that the boundary between the CL and the osteon was usually easier to detect than between the CL and the interstitial bone. The extracted mask was dilated with a round kernel of 2 pixels to include the CL, and the original mask of the osteon was subtracted from the dilated mask to obtain a 2-pixel (or 1.14 μm) thick circular layer representing the CL. Next, the bone around the CL was subdivided into several concentric layers having the same nominal thickness of the CL and located both inside and outside the osteon, in regions adjacent to the CL as well as further away (Figs. 1D and E). The layers were obtained by multiple cycles of erosion and dilation applied to the mask of the osteon (Figs. S1C and D). Specifically, we considered layers close to the CL and placed at 1 and 4 pixels (or 0.57 *µ*m and 2.28 μm) away from the CL border: 2 layers inside the osteon (I1, I2) and 2 layers outside (O1, O2). Two additional layers were defined 20 pixels (or 11.4 μm) away from the CL (I3 and O3, respectively). Inner layers include only osteonal bone, whereas outer layers may intersect with other (older) osteons and interstitial bone. Furthermore, all circular-shaped layers were radially sliced into 16 sectors (using MATLAB 2022a), which served to investigate the spatial variation of mineral content around the osteons (Fig. S1E). A custom MATLAB routine was implemented to compute the mean calcium content (Ca_Mean_), its standard deviation, as well as the frequency distribution of the calcium content, within each layer and in the whole osteon. All frequency distributions were smoothed with a running average filter (window size 10).

### Statistical Analysis

We compared differences in mineralization among the layers by performing a Kruskal-Wallis test, chosen due to the non-normal distribution of the data. When significant differences were detected, pairwise comparisons between layers were conducted using Tukey-Kramer’s post hoc test. We examined the strength of the relationships between the mean calcium content of osteon, CL and different layers using Spearman’s rank correlation, selected for its suitability with non-normal distribution of the data. Significance levels were set at p < 0.05. All statistical analyses were performed in MATLAB 2022a.

## Results

From a qualitative observation of the qBEI maps, the CL appears as a thin line around the osteon, usually somewhat brighter than the surrounding bone (Fig. 1E). In the SHG images there were no evident features allowing a clear discrimination between CL and adjacent lamellar bone (Fig. 1G). However, superimposing qBEI and SHG maps indicates that the CL tends to fall in regions with no (or very weak) SHG signal.

We explore possible links between the mineralization of the CL and immediately adjacent bone, considering regions both outside and inside the osteon, i.e. formed before and after CL deposition, respectively. We first report detailed findings on two osteons having low and high mean mineral content that are representative for younger and older tissue (assuming mineral content as an indirect measure of tissue age) and extracted from the adult (40 years old) and aged (81 years old) individual, respectively. We then present results of all 35 osteons, with mean calcium content ranging from 21.37 ± 2.26 wt % to 26.43 ± 2.2 wt %, and therefore covering a large spectrum of tissue age. Less mineralized younger osteons were more likely found in the adult individual, whereas older more mineralized osteons in the aged individual.

As a first example, we investigated an osteon with relatively low mean mineral content (Ca_Mean_ = 21.72 ± 2.09 wt %) coming from the adult individual and surrounded by older osteons and interstitial bone (Fig. 2A). SHG imaging reveals bone lamellae at the osteon periphery, confirming that the selected osteon is not interrupted by other osteons (Fig. 2B). The local and layer-based analysis of qBEI maps indicates a clear increase in mineral content of about 15% when going from the inner layers of the osteon to the CL, which has Ca_Mean_ of 25.54 ± 1.52 wt % (Fig. 2C). The difference in mineral content between the CL and the outer layers is much smaller (around 2%) but still statistically significant (p < 0.001). The same trend is confirmed when inspecting the frequency distributions of the calcium content of the layers, which essentially split into two groups (Fig. 2D): inner layers have similar distributions, overlapping with the distribution of the whole osteon, whereas the distributions of the outer layers are shifted towards higher mineral content. The frequency distribution of the CL displays an additional (albeit small) right-shift, making it distinguishable from all other curves. Owing to the circular shape of the osteon, the spatial correlation among the layers is further explored with a polar plot (Fig. 2E), based in our subdivision into 16 sectors (see Materials and Methods). A central observation is that in each sector, the mean calcium content of the CL follows very closely the mineral content of immediately adjacent bone located just outside the osteon and therefore deposited before the CL. This observation is quantified by an analysis of the Spearman correlation coefficient between the calcium content of the CL and of the adjacent outer layer O1 in the 16 sectors around osteon (Fig. 2F): with R = 0.9 (p < 0.001), such correlation can be considered as very strong^(30)^. Conversely, no spatial relationship between the mean mineral content of CL and inner layers is evident: the correlation between the calcium content of the CL and of the inner layer I1 in the 16 sectors is not statistically significant (Fig S2A, R = -0.4, p = 0.108).

**Fig. 2:**
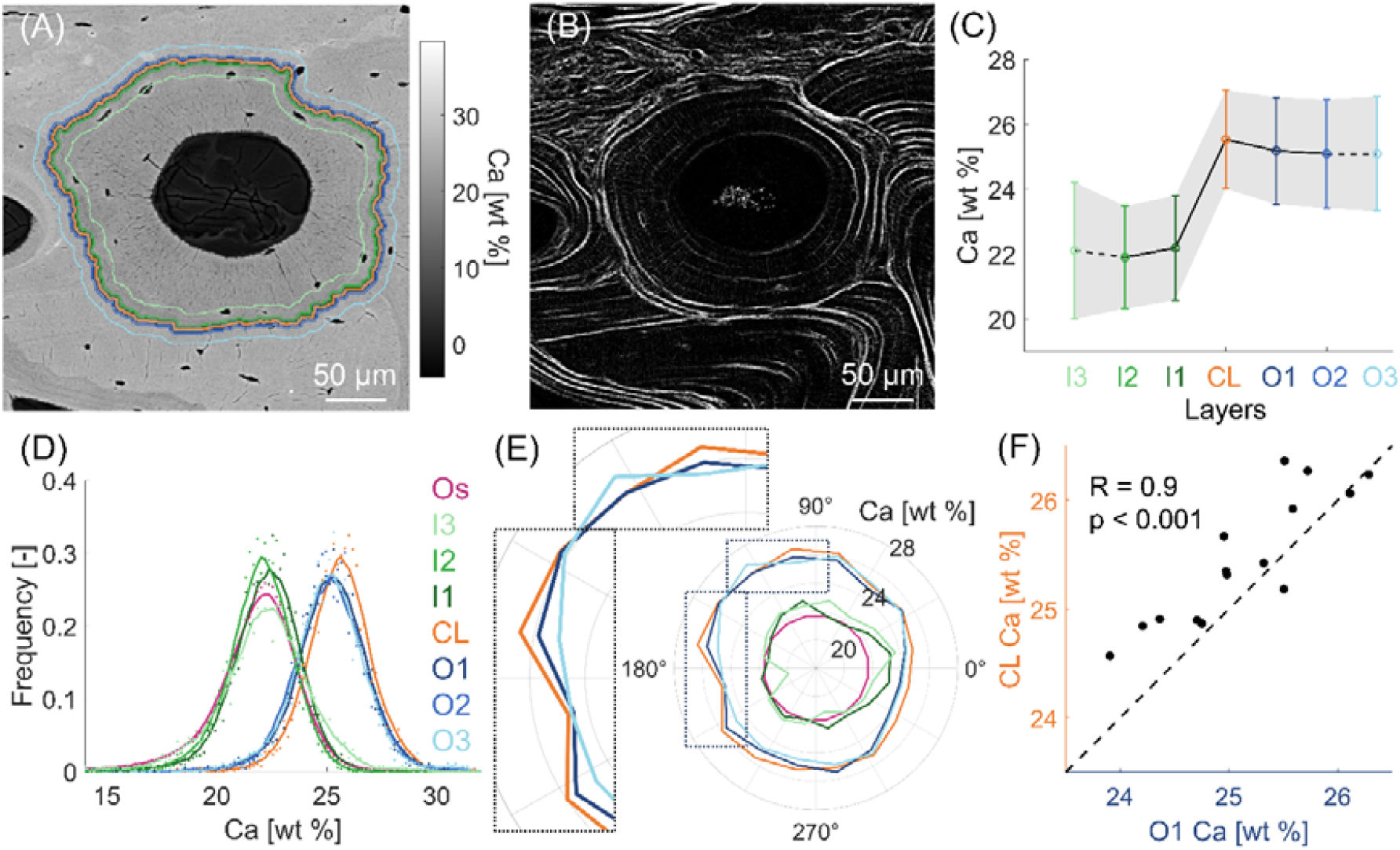
Mineral content analysis of a lowly mineralized younger osteon from the adult (40 years) bone sample. (A) qBEI maps with cement line (CL) and segmented concentric layers highlighted by the colored lines. (B) SHG image showing the lamellar organization of the uninterrupted osteon and neighboring bone. (C) Mean calcium content of the inner layers (I3, I2, I1), CL and outer layers (O1, O2, O3). Data shown as mean ± standard deviation (the shaded area delimits a one-standard deviation interval around the mean). Data points are connected by lines to visualize the trend: full lines connect points equally spaced, whereas dashed lines indicate that I3 and O3 are further away. (D) Frequency distributions of the calcium content of the entire osteon (Os), CL and segmented layers. Original data represented as scattered points and smoothed data (with moving average) as full lines. (E) Polar plot of the calcium content in selected regions (CL, O1, O3, I1, I3 and Os), averaged over 16 sectors around the osteon. In the polar plot the mean Ca content in each sector is plotted as distance from the center. (F) Correlation between the calcium content of the CL and of the adjacent outer layer (O1) considering all 16 sectors. The dotted line with a unit slope and passing through the origin is shown to improve the readability of the plot.

The second example is an osteon with higher mineral content (Ca_Mean_ = 25.95 ± 2.45 wt %) representative for older tissue and extracted from the aged individual. Again, this osteon is surrounded by a heterogeneous environment featuring older osteons as well as interstitial bone (Fig. 3A and B). The SHG analysis underlines a clear pattern of alternating and uninterrupted bone lamellae within the selected osteon (Fig. 3B). In this “aged” scenario, the CL has still higher mineral content than the osteon (about 9%) as well as than the external surrounding bone layers (about 5%, Fig. 3C). The frequency distributions of inner and outer layers are closer to each other’s w.r.t. the previous case of the younger osteon, whereas the distribution of the CL still displays a noticeable shift to higher mineral content (Fig. 3D). The polar plot suggests a statistically significant spatial relationship between the mineral content of CL and adjacent outer layers (Fig. 3E), which is characterized by a moderate correlation coefficient (R = 0.58, p < 0.05). Also in this example, the correlation between the CL and the inner layer I1 is not significant (Fig S2B, R = 0.41, p = 0.101).

**Fig. 3:**
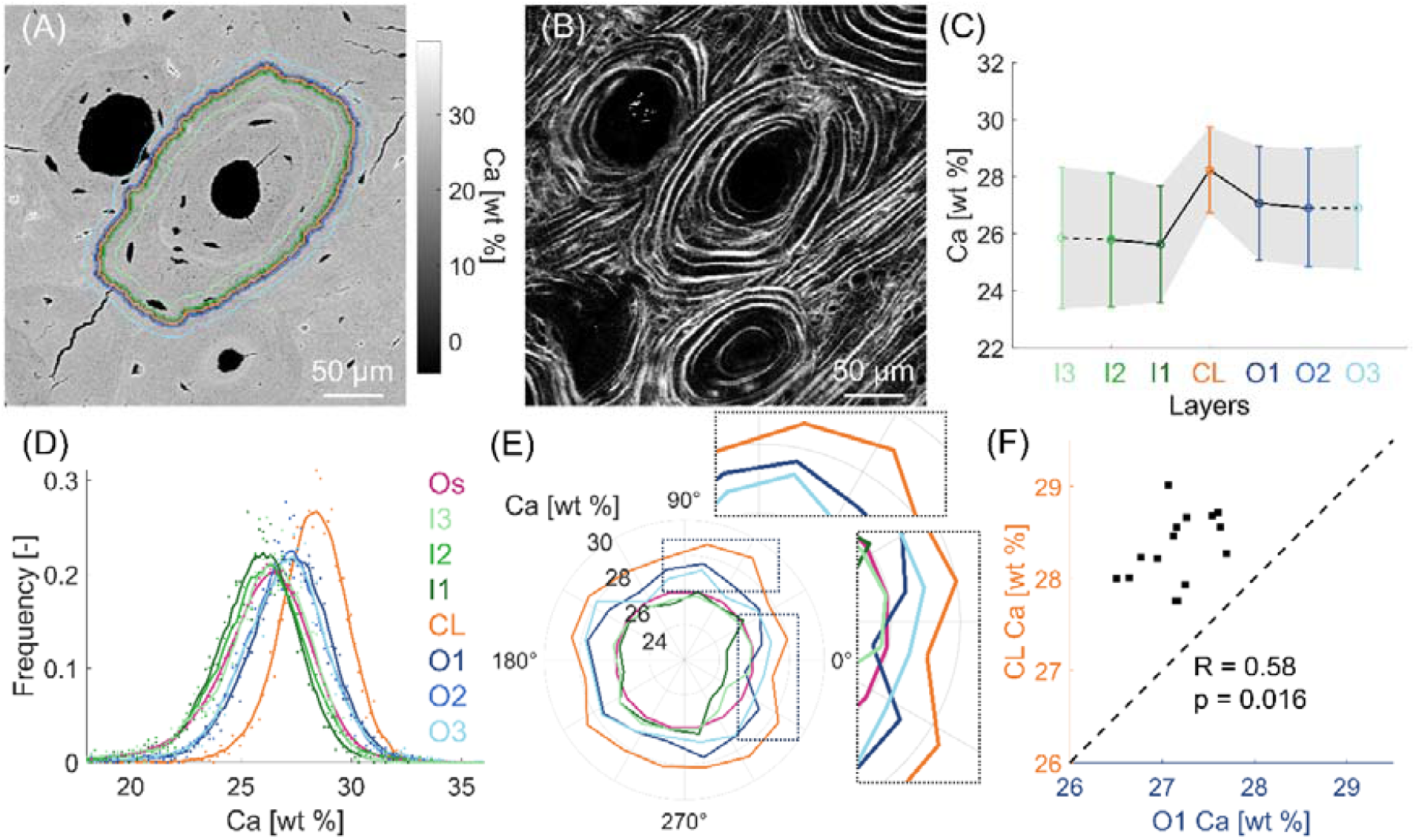
Mineral content analysis of a highly mineralized older osteon from the aged (81 years) bone sample. (A) qBEI maps with cement line (CL) and segmented concentric layers highlighted by the colored lines. (B) SHG image showing the lamellar organization. (C) Mean calcium content of the inner layers (I3, I2, I1), CL and outer layers (O1, O2, O3). Data shown as mean ± standard deviation (the shaded area delimits a one-standard deviation interval around the mean). Data points are connected by lines to visualize the trend: full lines connect points equally spaced, whereas dashed lines indicate that I3 and O3 are further away. (D) Frequency distributions of the calcium content of the entire osteon (Os), CL and segmented layers. (E) Polar plot of the calcium content in selected regions (CL, O1, O3, I1, I3 and Os), averaged over 16 sectors around the osteon. In the polar plot the mean Ca content in each sector is plotted as distance from the center. (F) Correlation between the calcium content of the CL and of the adjacent outer layer (O1) measured considering 16 discrete sectors around the osteon. The dotted line with a slope of one and passing through the origin is shown to improve the readability of the plot.

Extending the layer analysis to all osteons (Fig. 4), confirms that the CLs always exhibit the highest mean mineral content (27.35 ± 1.58 wt % Ca), not only in comparison with the layers inside the osteons but also with the layers outside. The difference in mineral content between the CL and the outer layers is less pronounced than between the CL and the inner layers. The outer layers show a smaller spreading in the mean mineral content when compared to the inner layers.

**Fig. 4:**
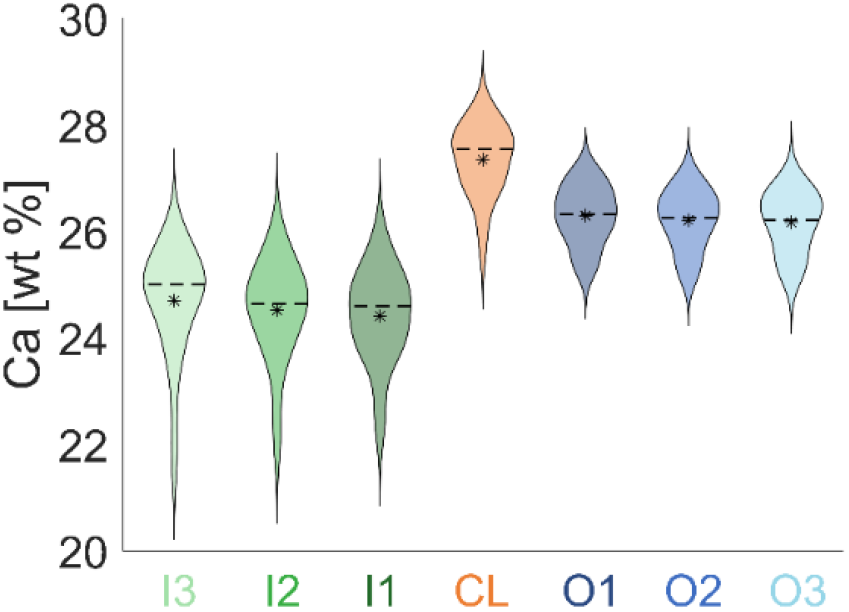
Violin plots (mirrored histograms) of the calcium content for each segmented region (CL, inner, and outer layers) considering all analyzed osteons (n=35). The mineral content of the CL is significantly higher (p < 0.001) than all other layers The violin plots are based on kernel density estimation. The median and mean values are indicated by a dashed line and an asterisk, respectively.

The mineralization of the CL is explored in Fig. 5. Firstly, we observed that the mineral content of the CL is always higher than the osteon mineral content, and that a strong correlation is present between them (R = 0.75, p < 0.001, Fig. 5A), suggesting that the CL mineral content progresses with tissue age. At the same time, the mineral content of the bone outside the osteon (represented by the outer layer O1) had only a moderate correlation (R = 0.4, p = 0.018) with the mineral content of the osteon (Fig. 5B). In contrast, the relationship between the mineral content of CL and outer layer O1 (Fig. 5C) displays a strong correlation (R = 0.75) with statistical significance (p < 0.001). As in Fig. 5A, all experimental points are above the line with a slope of 1 (i.e., 45°, dashed line in Figs. 5A-C). This confirms that the CL is the highest mineralized structure in all 35 investigated osteons compared to either the inside or the outside of the osteon. Virtually identical results are found when considering instead of the outer layer O1, the outer layer O2, which is further away from the CL (Figs. S3A and B). The strong correlation between the mineral content of the CL and the outer layer O1 motivates to investigate how the difference between these two quantities (CL – O1) behaves when plotted versus the osteon mineral content (Fig. 5D). The obtained correlation is strong (R = 0.75) and statistically significant (p < 0.001). The positive slope of the correlation (full line in Fig. 5D) shows that the mineral content of the CL deviates more from the mineral content of the surrounding, the older (i.e., the higher mineralized) the osteon is. An opposite trend is observed when plotting (CL – I1) versus the osteon mineral content (Fig. S8).

**Fig. 5:**
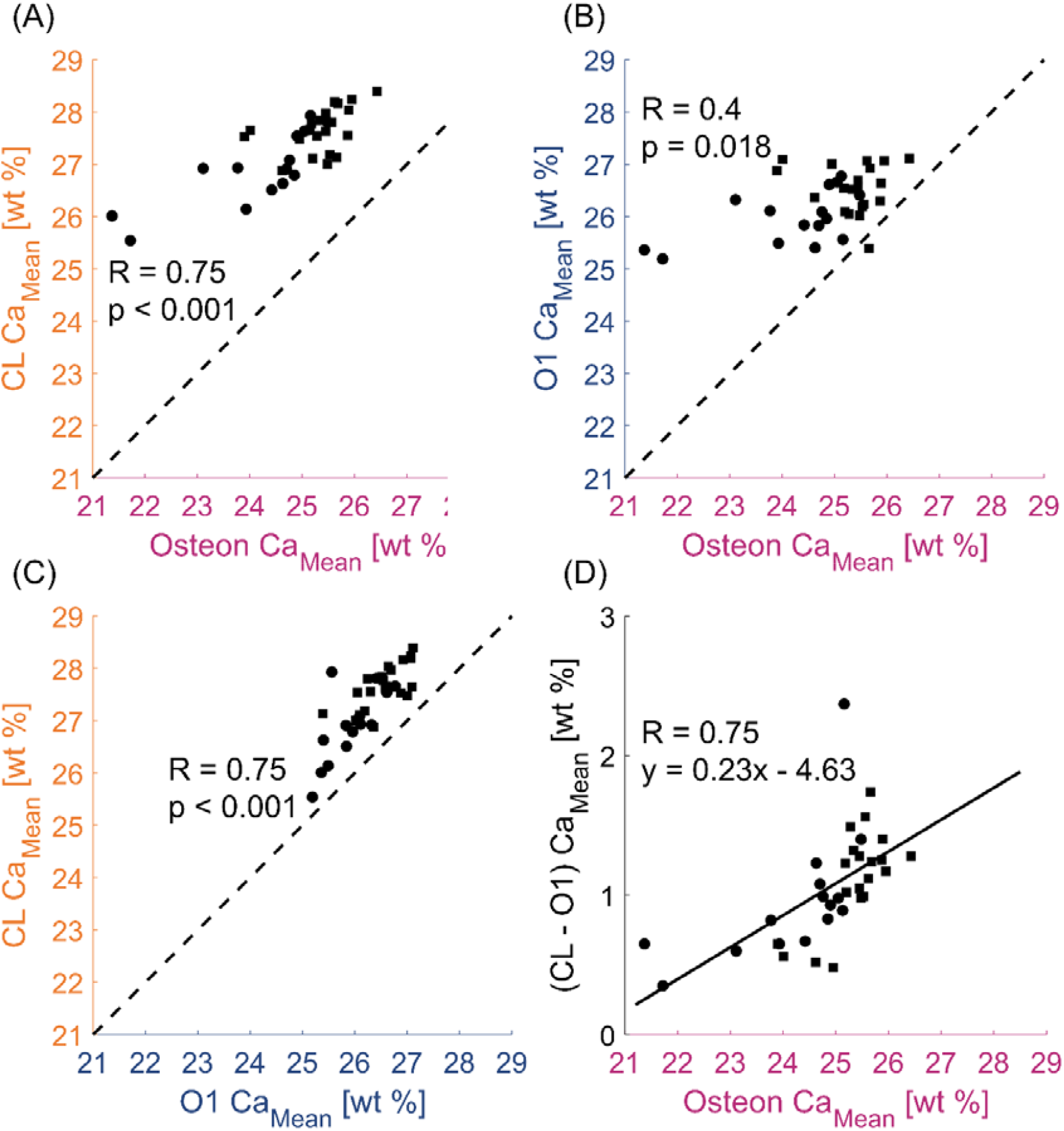
Mineral content of the CL in comparison to the mineral content of the osteon and the outer surrounding. (A) Correlation between average calcium content (Ca_Mean_) of the CL and the corresponding osteon, (B) outer layer O1 and osteon, (C) CL and O1, and (D) difference between CL and O1 (CL-O1) and osteon. The dotted line (with a unit slope and passing through the origin) is inserted to improve the readability of the plot. The full line in (D) represents the result of a linear regression. Circles represent osteons from the adult individual (40 years) and square from the aged individual (81 years).

Motivated by the strong correlation between the mineral content of the CL and of the outer layer O1 (Fig. 5C), we performed a spatially resolved analysis of the relationships between the mineralization of the CL and the outer/inner layers, by considering the calcium content within each circular sector around the osteon, for a total of 3920 different regions inspected (Fig. 6). Pooling all sector-wise data together (Fig. S5) confirmed the trend observed in Fig. 5, with CL and O1 showing the highest correlation and data points spread close to a line of unit slope. We applied the sector analysis (presented in Fig. 2F and 3F) to all osteons (Fig. 6A): for 21 out of 35 osteons, we detected a statistically significant correlation (p < 0.05) between the mineral content of the CL and outer layer O1, ranging from moderate (0.4 < R < 0.7) to strong (0.7 < R < 0.9). The osteons which did not show a significant correlation are likely to have higher mineral content (Ca_Mean_ > 24.6 wt %). The differences in mineral content between the CL and the closest outer layer (CL - O1) as well as between the CL and the closest inner layer (CL - I1) were then calculated in each discrete sector and averaged for each osteon (Fig. 6B). The results further underline that the degree of mineralization of the CL is extremely close to the mineral content of the outer layer O1, with the difference CL - O1 being always less than 2 wt % and showing a small increase for more mineralized osteons. In comparison, the differences between CL and I1 are higher (up to 3.5 - 4 wt %) and show a slight decreasing trend with osteon mineral content. A similar behavior is observed when considering all individual sectors (Fig. 6C). Here, we report results as 2D maps with differences (CL - O1) and (CL - I1) calculated in each sector but not averaged in the osteons. In Fig. 6C, the 16 individual sectors are on the vertical axis and the 35 osteons, sorted from lower to higher mineral content, are on the horizontal axis. Results indicate that even in discrete regions around the osteon, the mineral content of the CL is very similar to the degree of mineralization of the outer layer O1; this similarity being particularly strong for osteons with lower mineral content, i.e. assumed to be younger.

**Fig. 6.**
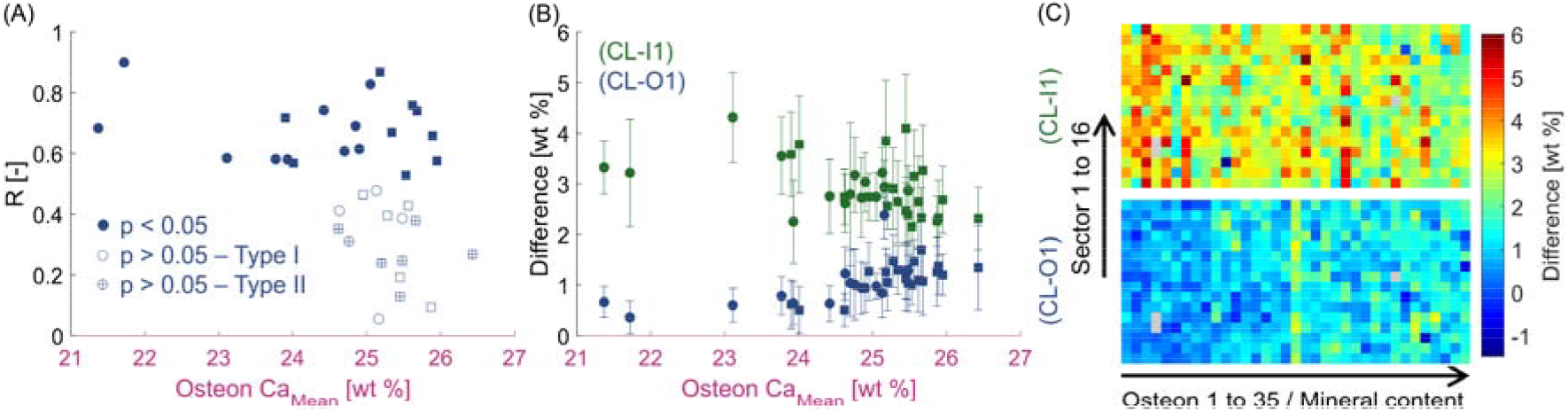
Spatial analysis of the mineral content. (A) Spearman correlation coefficients between the mineral content of the CL and the outer layer O1, computed according to a spatially resolved analysis around each osteon. All osteons that show a statistically significant correlation are of type I, whereas among the osteons where no correlation is detected, there are several osteons of type II (i.e., osteon-in-osteon). Circles represent osteons of the adult individual (40 years) and squares of the aged individual (80 years). (B) Differences in mineral content between the CL and the closest neighboring layers computed as average difference (CL-O1) and (CL-I1) in each circular sector around the osteon and plotted versus the mean degree of mineralization of the osteon. Data points represent the mean value and error bars the standard deviation for each osteon. Again, circles represent osteons from the adult individual (40 years) and square from the aged individual (81 years). (C) Local differences (CL-O1) and (CL-I1) calculated in each sector (vertical axis) and reported for each osteon sorted according to increasing mineral content (horizontal axis). A few grey pixels indicate sectors around the osteons where the mineral content could not be calculated, for example due to intersecting pores or cracks.

## Discussion

We investigated the mineral content of CL and neighboring tissue in human cortical bone using qBEI at submicrometer resolution, combined with a spatially resolved layer analysis. The main results of our analysis of 35 osteons from two individuals are summarized in Figures 5 and 6. An important assumption for the interpretation of the plots in these Figures is that the mineral content of the osteon serves as a proxy of tissue age. This allows us to deduce the following important aspects of CL mineralization.

Firstly, the obtained results are consistent for the osteons of both individuals despite their very different age (40 and 81 years, respectively). Consequently, we conclude that the tissue age (i.e., the mineral content of the osteon) is the crucial time parameter and is not confounded by the age of the individual. Secondly, the mineral content of the CL is always higher than of the corresponding osteon and always higher than the surrounding of the osteon. Thirdly, the strong correlation between the mineral content of the CL and the mineral content of the osteon (Fig. 5A) demonstrates that also the mineral content of the CL evolves in time following a characteristic time course, with differences with respect to the mineralization of the osteon. This is in agreement with previous findings on postmenopausal women^(15)^. To describe the mineralization kinetics of the CL, a separation into a fast primary mineralization and a much slower secondary mineralization is useful. Osteon and CL are deposited within the same remodeling cycle and therefore have a similar tissue age. Nevertheless, the mineral content of the CL is always higher than the corresponding osteons, even for relatively young osteons. This suggests that in primary mineralization, the CL can incorporate more mineral than the osteon. Fourthly, the mineral content of the CL after primary mineralization is closely related to the mineral content of the surrounding in which the CL and the new osteon are formed. This is true not only when considering the average mineralization (Fig. 5C) but extends to the very local environment of the CL as our analysis shows when the CL is subdivided into 16 segments (Fig. 6). This indicates that the mineral content of the CL depends, in addition to osteon age (Fig. 5A), also on the degree of mineralization of the region where the CL is formed. The local influence of the environment on the initial mineral content of the CL could be explained considering the events leading to CL formation. The common picture is that the CL is formed on the eroded surface immediately after, or perhaps even during, bone resorption. In this process, osteoclasts generate a local acidic environment allowing to mobilize the inorganic phase of bone^(31)^. Mineral ions/clusters are then transported through the osteoclasts and released nearby in the extracellular space^(32)^. At the same time, osteoclasts (and osteoprogenitors cells) produce osteopontin, a highly charged and phosphorylated protein, well-known for its high affinity to bind calcium ions^(33)^. The presence of osteopontin as one of the first molecules deposited during CL formation, together with its ability to bind calcium ions and mineral crystals, could enhance the accumulation of mineral in the newly forming CL^(34)^. The higher the mineral content of the region undergoing resorption, the more mineral ions/clusters may be released by the osteoclasts and then locally “captured” through osteopontin to build the CL. Such local “recycling” of mineral may affect the primary mineralization of the CL, which may be faster in the presence of more mineral ions and could explain the strong correlation between the mineral content of the CL and of the bone outside the osteon (layer O1, Fig. 5C). This correlation is higher than the correlation between osteon and O1 (Fig. 5B), meaning that secondary mineralization alone cannot be the only cause for the observed trend. The formation of the bone inside the osteon is somewhat “dephased”, both in space and time, with respect to the deposition of the CL as a critical density of osteoblast-lineage cells is required^(1)^. In osteonal remodeling, the distance separating the initial resorption and formation fronts can be even around 1 mm^(1)^: assuming that osteoclasts resorb bone with a speed of at about 20-40 *µ*m/day^(35)^, this distance would correspond to 20-50 days. Thus, the concentration of mineral ions from resorbed bone available during osteonal bone deposition may be different than during the formation of the CL. This could be one reason why there is only a weak correlation (in several cases not statistically significant) between CLs and bone layers inside the osteon, particularly when data are averaged in the discrete sectors around the osteon (Fig. S7).

Fifthly, Figure 5D shows that the difference between the mineral content of the CL and of its outside surrounding increases with tissue age. Since the surrounding of the osteon is old bone tissue, which is already in advanced secondary mineralization, it can be assumed that its mineral content hardly changes any more. Consequently, the increase in mineral content corresponds to the rather slow secondary mineralization of the CL. We may even attempt to estimate the rate of CL secondary mineralization by looking at Fig. 5D. Based on available data on turnover rate in osteonal bone^(36–38)^, we estimate an age difference of about 15 years between the youngest and the oldest osteon. This means that the mineral content of the CL increases due to secondary mineralization of about 2 wt % over a course of 15 years. In contrast, within the same time span, the increase in osteon mineral content is much higher and about 5 wt % (Fig. 5A). A secondary mineralization, which is slower in the CL compared to the osteon, is also consistent with the finding that the higher the mean mineral content of the osteon, the less is the difference in mineralization between the osteon and the CL (Fig. S4).

Sixthly, we could identify a reason why the correlation between the mineral content of the CL and the outer surrounding is weaker. Many of these osteons with a weak correlation are classified as type II osteons^(39–41)^ also known as “osteon-in-osteon”^(42)^. This term refers to a specific type of relatively large osteon containing a smaller osteon in its center, both being surrounded by concentric CL^(39)^. In type II osteons, the degree of mineralization of the external environment around the osteon tends to be more homogenous in comparison to osteons surrounded by both interstitial bone and older osteons (Fig. S6). The frequency of type II osteons increases with age^(39,43)^, although their precise function and significance in bone remain largely unknown. In our dataset “osteon-in-osteon” are more likely seen in the bone sample of the aged donor. In these structures, only the CL of the inner osteon was considered as the outer osteon was in some cases interrupted by other osteons.

In summary, both the osteon and the CL undergo a mineralization process, which lasts over several years. The kinetics of this process can be described by a very fast primary mineralization followed by a much slower secondary mineralization. For bone this is well understood and can be even quantitatively described^(17)^. Here, we provide information on the mineralization kinetics of the CL. After primary mineralization, the mineral content of the CL is substantially higher than the mineral content in osteonal bone, possibly beyond the content of fully mineralized bone. Most importantly, the mineral content of the CL after primary mineralization is clearly not universal but depends on the location where the remodeling event occurs, with the CL being more mineralized when the osteon is surrounded by bone of higher mineral content. This could be due to a local recycling of mineral: more ions released during bone resorption allows for a more mineralized CL. The CL shows also a secondary mineralization which proceeds at much slower rates than in osteonal bone. One reason for the slower pace may be the high mineral content already attained at the end of primary mineralization. The interplay between primary and secondary mineralization of the CL can be explored with the help of a previously developed mathematical model for the mineralization process of bone^(17)^ (Supplementary Information and Fig. S9): if the CL has an augmented primary mineralization (which is also location-specific), the secondary mineralization needs to be reduced to avoid attaining unrealistically high values of the mineral content (Fig. S9). For an intuitive representation of CL mineralization, one could imagine that the “driving force” to mineralize is somehow proportional to the difference to the maximum mineral content from the actual mineral content. As the actual mineral content increases (due to mineralization), there is a natural decrease in the “driving force”, leading to an asymptotic approach of the mineral content to its maximum value. This is exactly how a simple thermostat works: the heat flux which is regulated through the thermostat is proportional to the difference of the actual temperature to the set point temperature.

The found higher mineral content of all CLs compared to the mineral content of the neighboring osteonal and interstitial bone (Figs. 4 and 5A, C) agrees with previous works based on qBEI^(15)^, X-ray phase contrast nanotomography^(10)^, and energy dispersive X-ray spectroscopy^(13)^. One possible explanation for the observed higher mineral content of the CL has been obtained by analyzing osteonal bone with focused ion beam scanning electron microscopy (FIB-SEM). A network of nanochannel, much smaller than the canaliculi connecting osteocytes lacuna has been found in osteonal bone but much less in the CLs bordering osteons^(25)^. The high mineral content of the CL measured with qBEI (as well as with other techniques having a submicrometer resolution), may be due to a reduced nanoporosity inside the probed volume, rather than to a higher degree of mineralization of the organic matrix^(25)^. The origin of nanochannels in bone tissue is still not understood. One possible explanation is that they form during tissue maturation and can be considered as leftovers of growing mineral domains^(25)^. As the CL reaches a very high mineral content already at the end of the primary mineralization, the slower secondary mineralization may be due to the closing of the nanochannels.

The following limitations of this work should be mentioned. First, we acquired qBEI maps with a resolution of 0.57 μm/pixel and the typical width of the CL in 2D sections is about 1-3 *µ*m, so there is a fraction of pixels only partially filled by the CL, impacting the detected mineral content. A higher resolution would allow for an improved characterization of the CL; however, at higher magnifications the backscattered electrons signal is not only influenced by the local calcium content but also by other features such as the arrangement of the mineralized collagen fibrils an the crystallographic orientations ^(25)^. Second, our samples were obtained postmortem which did not allow the use of fluorescent labels that are the only possibility to precisely determine tissue age, to assess the time course of mineralization in newly formed bone^(16)^, and perhaps even to investigate possible differences in early stages of mineralization between CL and osteonal bone. Therefore, in our work as well as in the literature^(15,24,44,45)^, the osteon mineral content is used as an indicator of tissue age, but it cannot be excluded that different osteons mineralize with different kinetics. Third, we used manual contouring to identify the outer border of the osteons based on different gray values between osteon and CL. This is a time-consuming approach which limits the number of osteons that could be investigated in a reasonable timeframe. Using an algorithmic approach may provide a rapid identification of the CLs allowing to include more samples and hundreds of osteons coming from individuals with different ages and considering additional anatomical locations. Fourth, the qBEI maps of the mineral content and the SHG images showing fiber organization could not be superimposed with enough accuracy to be able to unambiguously locate the SHG signal directly at the CL: considering the very tiny dimension of the CL, already small deviations from the CL location can lead to high inaccuracies.

In conclusion, we showed a close spatial interplay between the mineralization of the CL and the mineral content of the surrounding older bone, suggesting that CL mineralization is different in different osteons. The fact that the process of bone mineralization is not universal as previously believed but depends on anatomical sites has recently been demonstrated by analyzing mineral characteristics at modeling and remodeling surfaces and may be linked to different avenues to transport mineral ions to the mineralization sites ^(24)^. Here, we extend this novel concept by providing evidence that, even within the same anatomical location and remodeling cycle, there are differences in bone mineralization between the CL and the subsequently formed osteonal bone, which may be also due to dissimilar concentrations of mineral ions depending on the very local mineral content of the resorbed regions. Bone diseases or treatments affecting bone mineralization may have a direct impact on the mineral content, and therefore on the mechanical behavior of the CLs.

## Supporting information

See Supplementary Information

## Conflicts of Interest

The authors declare no competing interests.

## Acknowledgements

AC is supported by the French Community of Belgium as part of a FRIA (Fund for Research Training in Industry and Agriculture) grant (n°1E03621F). We warmly thank Phaedra Messmer, Petra Keplinger, and Sonja Lueger from LBIO for the excellent sample preparation. MH and SB gratefully acknowledge financial support from the Austrian Social Health Insurance Fund (OEGK) and the Austrian Workers’ Compensation Board (AUVA). MR and RW thank the Max Planck Queensland Centre for the Materials Science of Extracellular Matrices for support.

## Notes

### Competing Interest Statement

The authors have declared no competing interest.

## References

1. Lassen NE, Andersen TL, Pløen GG, Søe K, Hauge EM, Harving S, Eschen GET, Delaisse J. Coupling of Bone Resorption and Formation in Real Time: New Knowledge Gained From Human Haversian BMUs. J. Bone Miner. Res. 2017 Jul;32(7):1395–405.

2. Lamarche BA, Thomsen JS, Andreasen CM, Lievers WB, Andersen TL. 2D size of trabecular bone structure units (BSU) correlate more strongly with 3D architectural parameters than age in human vertebrae. Bone. 2022 Jul;160:116399.

3. Tits A, Blouin S, Rummler M, Kaux J-F, Drion P, Van Lenthe GH, Weinkamer R, Hartmann MA, Ruffoni D. Structural and functional heterogeneity of mineralized fibrocartilage at the Achilles tendon-bone insertion. Acta Biomater. 2023 Aug;166:409–18.

4. Gupta HS, Schratter S, Tesch W, Roschger P, Berzlanovich A, Schoeberl T, Klaushofer K, Fratzl P. Two different correlations between nanoindentation modulus and mineral content in the bone–cartilage interface. J. Struct. Biol. 2005 Feb;149(2):138–48.

5. Nalla RK, Kruzic JJ, Kinney JH, Ritchie RO. Mechanistic aspects of fracture and R-curve behavior in human cortical bone. Biomaterials. 2005 Jan;26(2):217–31.

6. Koester KJ, Ager JW, Ritchie RO. The true toughness of human cortical bone measured with realistically short cracks. Nat. Mater. 2008 Aug;7(8):672–7.

7. Mohsin S, O’Brien FJ, Lee TC. Osteonal crack barriers in ovine compact bone. J. Anat. 2006 Jan;208(1):81–9.

8. Zimmermann EA, Gludovatz B, Schaible E, Busse B, Ritchie RO. Fracture resistance of human cortical bone across multiple length-scales at physiological strain rates. Biomaterials. 2014 Jul;35(21):5472–81.

9. Wagermaier W, Klaushofer K, Fratzl P. Fragility of Bone Material Controlled by Internal Interfaces. Calcif. Tissue Int. 2015 Sep;97(3):201–12.

10. Gauthier R, Follet H, Olivier C, Mitton D, Peyrin F. 3D analysis of the osteonal and interstitial tissue in human radii cortical bone. Bone. 2019 Oct;127:526–36.

11. Gustafsson A, Wallin M, Khayyeri H, Isaksson H. Crack propagation in cortical bone is affected by the characteristics of the cement line: a parameter study using an XFEM interface damage model. Biomech. Model. Mechanobiol. 2019 Aug;18(4):1247–61.

12. Burr DB, Schaffler MB, Frederickson RG. Composition of the cement line and its possible mechanical role as a local interface in human compact bone. J. Biomech. 1988 Jan 1;21(11):939–45.

13. Skedros JG, Holmes JL, Vajda EG, Bloebaum RD. Cement lines of secondary osteons in human bone are not mineral-deficient: New data in a historical perspective. Anat. Rec. A. Discov. Mol. Cell. Evol. Biol. 2005 Sep;286A(1):781–803.

14. Langer M, Pacureanu A, Suhonen H, Grimal Q, Cloetens P, Peyrin F. X-Ray Phase Nanotomography Resolves the 3D Human Bone Ultrastructure. Zuo X-N, editor. PLoS ONE. 2012 Aug 29;7(8):e35691.

15. Milovanovic P, vom Scheidt A, Mletzko K, Sarau G, Püschel K, Djuric M, Amling M, Christiansen S, Busse B. Bone tissue aging affects mineralization of cement lines. Bone. 2018 May;110:187–93.

16. Roschger P, Paschalis EP, Fratzl P, Klaushofer K. Bone mineralization density distribution in health and disease. Bone. 2008 Mar;42(3):456–66.

17. Ruffoni D, Fratzl P, Roschger P, Klaushofer K, Weinkamer R. The bone mineralization density distribution as a fingerprint of the mineralization process. Bone. 2007 May;40(5):1308–19.

18. Bala Y, Farlay D, Delmas PD, Meunier PJ, Boivin G. Time sequence of secondary mineralization and microhardness in cortical and cancellous bone from ewes. Bone. 2010 Apr;46(4):1204–12.

19. Lukas C, Ruffoni D, Lambers FM, Schulte FA, Kuhn G, Kollmannsberger P, Weinkamer R, Müller R. Mineralization kinetics in murine trabecular bone quantified by time-lapsed in vivo micro-computed tomography. Bone. 2013 Sep;56(1):55–60.

20. McKEE MD, Nanci A. Osteopontin and the Bone Remodeling Sequence: Colloidal-Gold Immunocytochemistry of an Interfacial Extracellular Matrix Protein a. Ann. N. Y. Acad. Sci. 1995 Aug;760(1):177–89.

21. Everts V, Delaissé JM, Korper W, Jansen DC, Tigchelaar-Gutter W, Saftig P, Beertsen W. The Bone Lining Cell: Its Role in Cleaning Howship’s Lacunae and Initiating Bone Formation. J. Bone Miner. Res. 2002 Jan 1;17(1):77–90.

22. Dr. McKee MD, Farach-Carson MC, Butler WT, Hauschka PV, Nanci A. Ultrastructural immunolocalization of noncollagenous (osteopontin and osteocalcin) and plasma (albumin and α2HS-glycoprotein) proteins in rat bone. J. Bone Miner. Res. 1993 Apr 1;8(4):485–96.

23. Sims NA, Martin TJ. Coupling the activities of bone formation and resorption: a multitude of signals within the basic multicellular unit. BoneKEy Rep. [Internet]. 2014 Jan 8 [cited 2024 Mar 29];3. Available from: http://www.portico.org/Portico/article?article=pgk2ph97srv

24. Roschger A, Wagermaier W, Gamsjaeger S, Hassler N, Schmidt I, Blouin S, Berzlanovich A, Gruber GM, Weinkamer R, Roschger P, Paschalis EP, Klaushofer K, Fratzl P. Newly formed and remodeled human bone exhibits differences in the mineralization process. Acta Biomater. 2020 Mar;104:221–30.

25. Tang T, Landis W, Blouin S, Bertinetti L, Hartmann MA, Berzlanovich A, Weinkamer R, Wagermaier W, Fratzl P. Subcanalicular Nanochannel Volume Is Inversely Correlated With Calcium Content in Human Cortical Bone. J. Bone Miner. Res. 2022 Dec 14;jbmr.4753.

26. Roschger P, Fratzl P, Eschberger J, Klaushofer K. Validation of quantitative backscattered electron imaging for the measurement of mineral density distribution in human bone biopsies. Bone. 1998 Oct;23(4):319–26.

27. Roschger P, Eschberger J Jr HP. Formation of Ultracracks in Methacrylate-Embedded Undecalcified Bone Samples by Exposure to Aqueous Solutions.

28. Lukas C, Kollmannsberger P, Ruffoni D, Roschger P, Fratzl P, Weinkamer R. The Heterogeneous Mineral Content of Bone—Using Stochastic Arguments and Simulations to Overcome Experimental Limitations. J. Stat. Phys. 2011 Jul;144(2):316–31.

29. Hartmann MA, Blouin S, Misof BM, Fratzl-Zelman N, Roschger P, Berzlanovich A, Gruber GM, Brugger PC, Zwerina J, Fratzl P. Quantitative Backscattered Electron Imaging of Bone Using a Thermionic or a Field Emission Electron Source. Calcif. Tissue Int. 2021 Aug;109(2):190–202.

30. Hinkle DE, Wiersma W, Jurs SG. Applied statistics for the behavioral sciences [Internet]. 5th ed. Boston, Mass., [London]: Houghton MifflinC; [Hi Marketing] (distributor); 2003. Available from: http://catalog.hathitrust.org/api/volumes/oclc/50716608.html

31. Teitelbaum SL. Bone Resorption by Osteoclasts. Science. 2000 Sep;289(5484):1504–8.

32. Vääräniemi J, Halleen JM, Kaarlonen K, Ylipahkala H, Alatalo SL, Andersson G, Kaija H, Vihko P, Väänänen HK. Intracellular Machinery for Matrix Degradation in Bone-Resorbing Osteoclasts. J. Bone Miner. Res. 2004 Sep 1;19(9):1432–40.

33. McKee MD, Nanci A. Osteopontin at mineralized tissue interfaces in bone, teeth, and osseointegrated implants: Ultrastructural distribution and implications for mineralized tissue formation, turnover, and repair. Microsc. Res. Tech. 1996 Feb 1;33(2):141–64.

34. Tang T, Landis W, Raguin E, Werner P, Bertinetti L, Dean M, Wagermaier W, Fratzl P. A 3D Network of Nanochannels for Possible Ion and Molecule Transit in Mineralizing Bone and Cartilage. Adv. NanoBiomed Res. 2022 Apr 28;2100162.

35. Webster J. Integrated Bone Tissue Physiology: Anatomy and Physiology. 2001;

36. Eriksen EF, Melsen F, Sod E, Barton I, Chines A. Effects of long-term risedronate on bone quality and bone turnover in women with postmenopausal osteoporosis. Bone. 2002 Nov;31(5):620–5.

37. Parfitt AM. Misconceptions (2): turnover is always higher in cancellous than in cortical bone. Bone. 2002 Jun;30(6):807–9.

38. Lerebours C, Weinkamer R, Roschger A, Buenzli PR. Mineral density differences between femoral cortical bone and trabecular bone are not explained by turnover rate alone. Bone Rep. 2020 Dec;13:100731.

39. Ericksen M. Histologic estimation of age at death using the anterior cortex of the femur. J. Phys. Anthropol. 1991 Dec;84(2):171–9.

40. Maggiano IS, Maggiano CM, Clement JG, Thomas CDL, Carter Y, Cooper DML. Three-dimensional reconstruction of Haversian systems in human cortical bone using synchrotron radiation-based micro-CT: morphology and quantification of branching and transverse connections across age. J. Anat. 2016 May;228(5):719–32.

41. van Tol AF, Roschger A, Repp F, Chen J, Roschger P, Berzlanovich A, Gruber GM, Fratzl P, Weinkamer R. Network architecture strongly influences the fluid flow pattern through the lacunocanalicular network in human osteons. Biomech. Model. Mechanobiol. 2020 Jun;19(3):823–40.

42. Redelstorff R. Unique bone histology in partial large bone shafts from Aust Cliff (England, Upper Triassic): an early independent experiment in gigantism. Acta Palaeontol. Pol. [Internet]. 2014 [cited 2024 Sep 18]; Available from: http://www.app.pan.pl/article/item/app20120073.html

43. Yoshino M, Imaizumi K, Miyasaka S, Seta S. Histological estimation of age at death using microradiographs of humeral compact bone. Forensic Sci. Int. 1994;64(2):191–8.

44. Gamsjaeger S, Hofstetter B, Zwettler E, Recker R, Gasser JA, Eriksen EF, Klaushofer K, Paschalis EP. Effects of 3 years treatment with once-yearly zoledronic acid on the kinetics of bone matrix maturation in osteoporotic patients. Osteoporos. Int. 2013 Jan;24(1):339–47.

45. Gourion-Arsiquaud S, Burket JC, Havill LM, DiCarlo E, Doty SB, Mendelsohn R, Van Der Meulen MC, Boskey AL. Spatial Variation in Osteonal Bone Properties Relative to Tissue and Animal Age. J. Bone Miner. Res. 2009 Jul 1;24(7):1271–81.

