## Supplementary material for "The mineralization of osteonal cement line depends on where the osteon is formed": See Supplementary Information

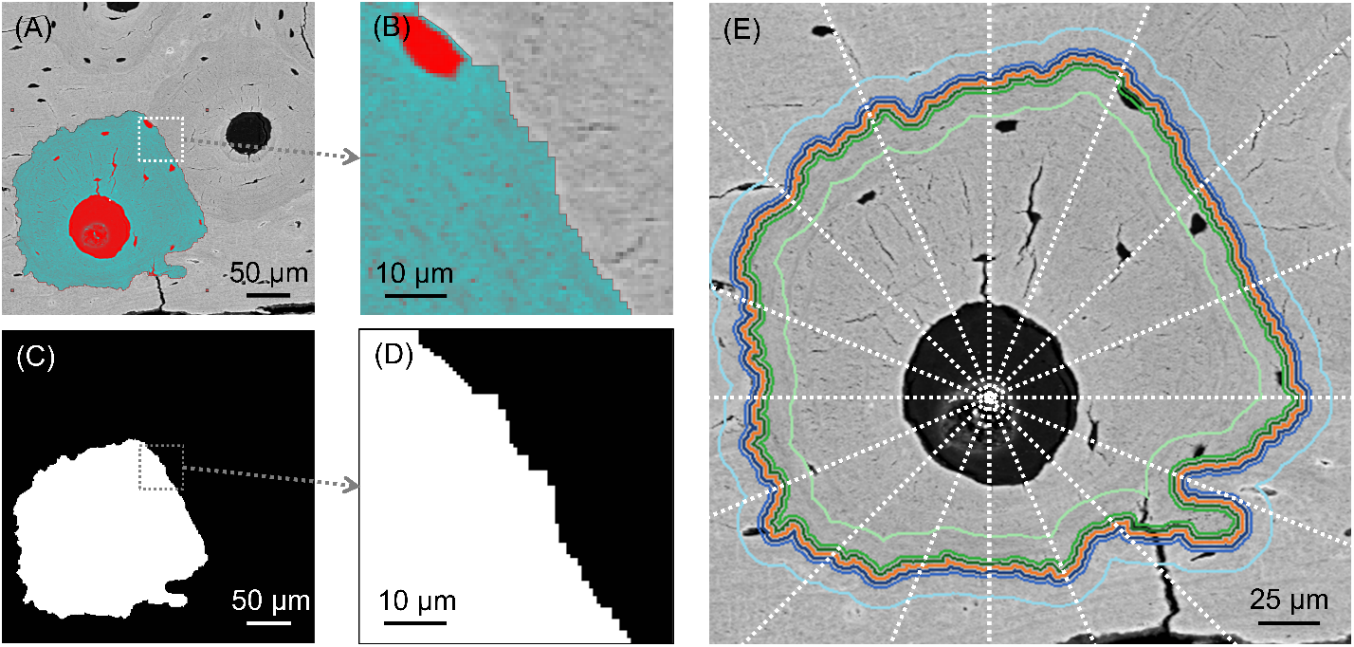

Fig. S1. Layer analysis of human osteonal bone. (A) Mask of the osteon obtained by manual segmentation of the quantitative backscattered electron imaging (qBEI) maps and (B) magnified view of the osteon border delimited by the CL, appearing brighter in qBEI. (C) Binarized mask of the osteon used to obtain each layer and (D) magnified view of the osteon border. (E) Subdivision of the circular-shaped layers into 16 sectors to investigate the spatial variation of mineral content around the osteons.

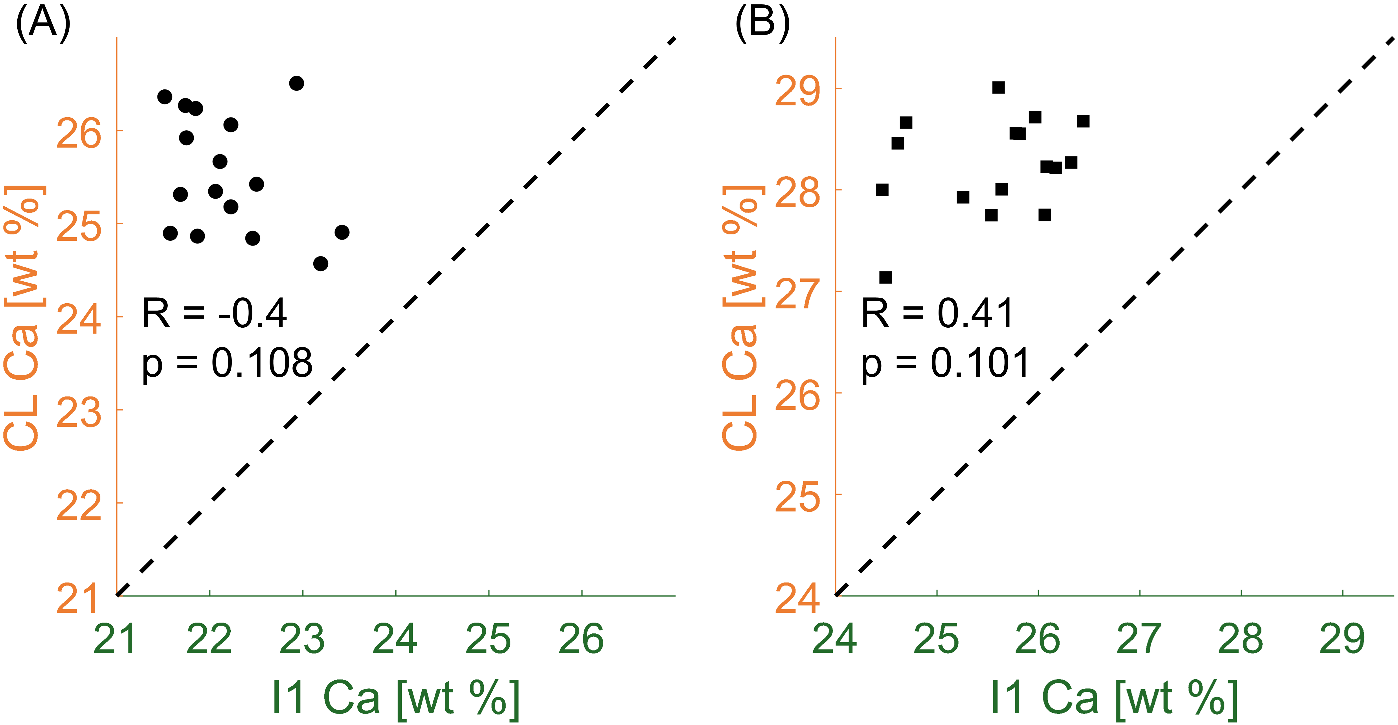

Fig. S2. Correlation between the calcium content of the CL and of the adjacent inner layer (I1) measured in 16 discrete sectors around the osteon. Results are shown for (A) an osteon with a low mineral content (Ca_Mean_ = 21.72 ± 2.09 wt %) and (B) an osteon with a high mineral content (Ca_Mean_ = 25.95 ± 2.45 wt %). The dotted line with a unit slope and passing through the origin is shown to improve the readability of the plot. Circles represent osteons from the adult individual (40 years) and square from the aged individual (81 years).

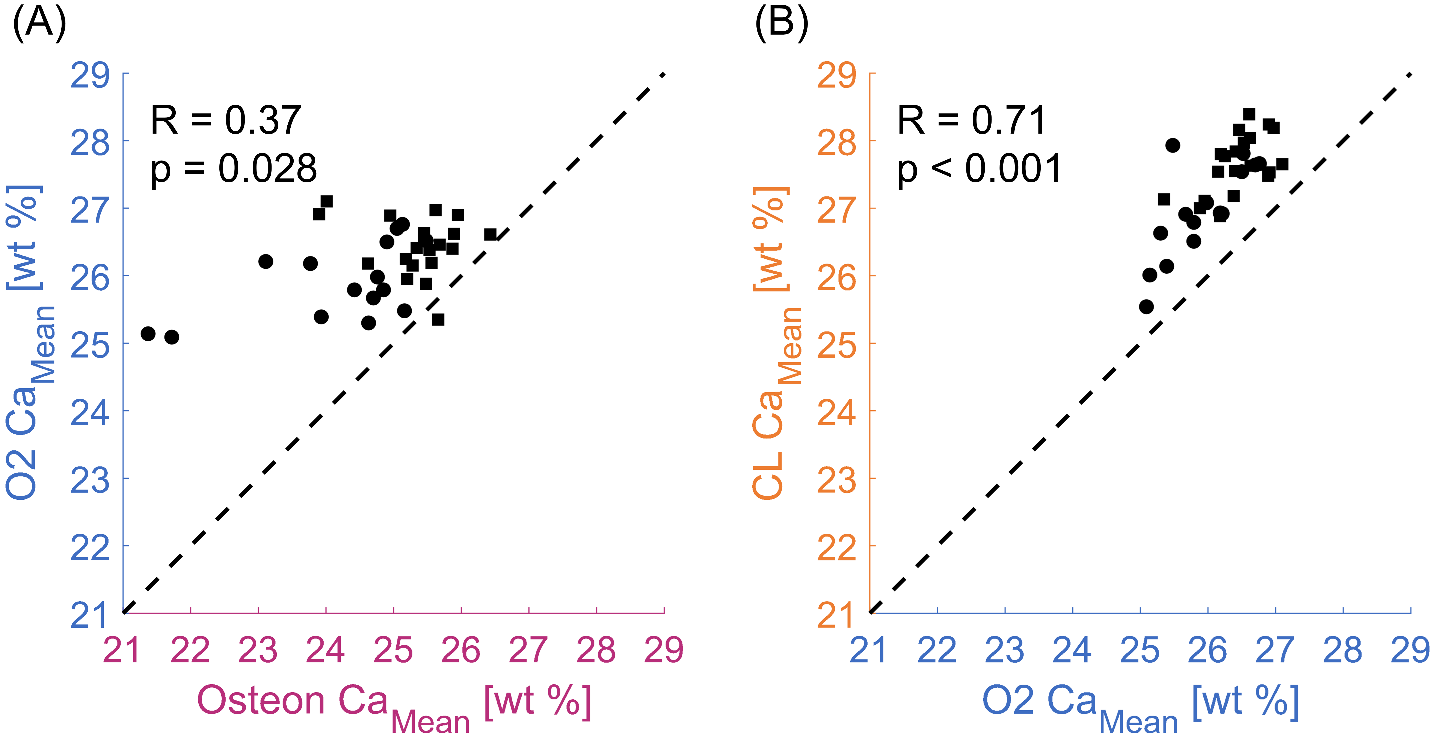

Fig. S3. Mineral content of osteon, CL and outer region. Correlation between the average calcium content (Ca_Mean_) of (A) the osteon and the outer layer O2, and (B) the CL and O2. The dotted line with a unit slope and passing through the origin is shown to improve the readability of the plot. Circles represent osteons from the adult individual (40 years) and square from the aged individual (81 years).

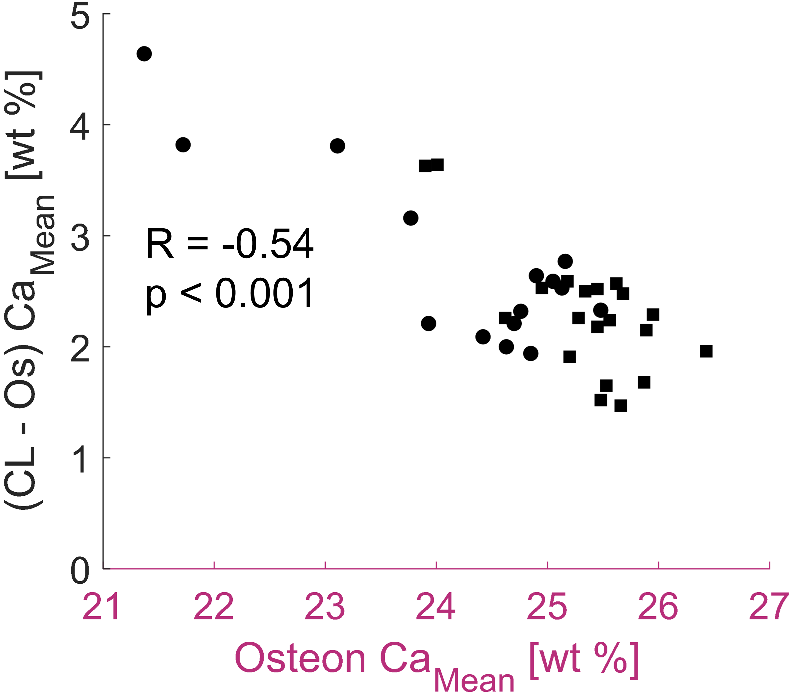

Fig. S4. Differences in the mean degree of mineralization between CL and corresponding osteon plotted versus the average calcium content of the osteon as a surrogate measure of osteon age. Circles represent osteons from the adult individual (40 years) and square from the aged individual (81 years).

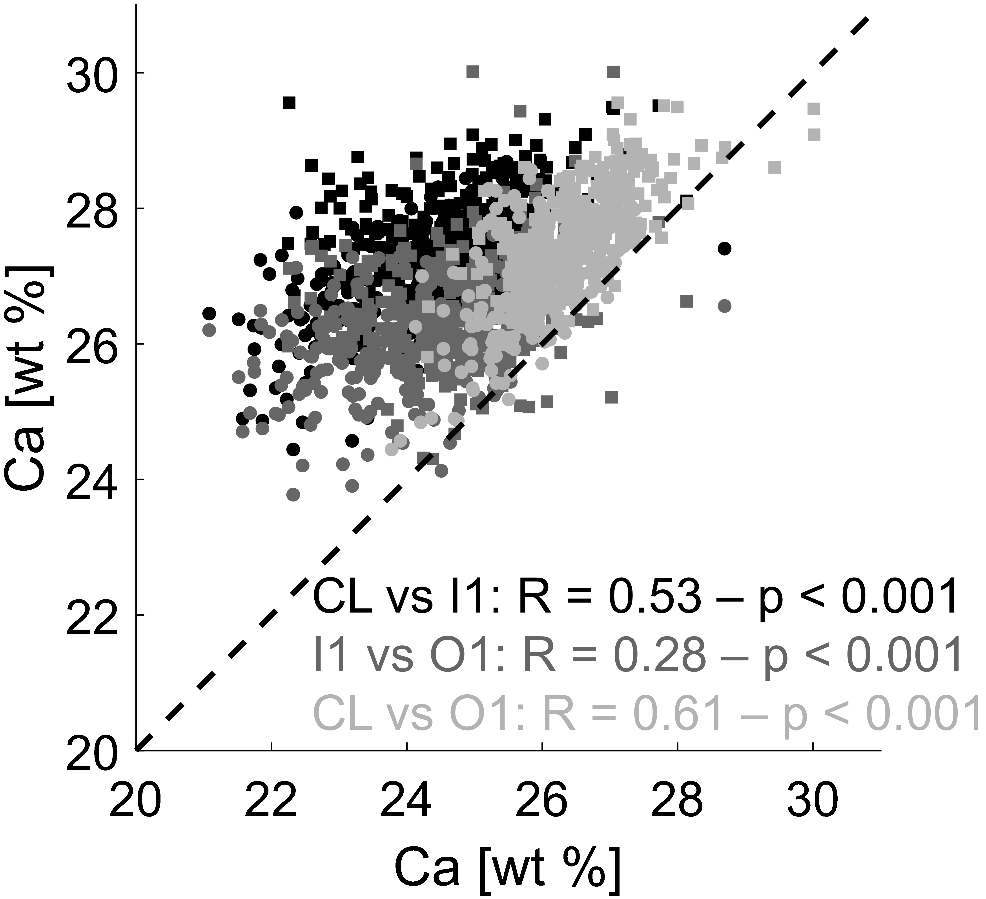

Fig. S5. Correlations between the calcium content of the CL and adjacent layers with datapoints representing individual sectors. The following regions are correlated: CL versus inner layer I1, CL versus outer layer O1 and inner (I1) versus outer layer (O1). The dotted line with a unit slope and passing through the origin is shown to improve the readability of the plot. Circles represent osteons from the adult individual (40 years) and square from the aged individual (81 years).

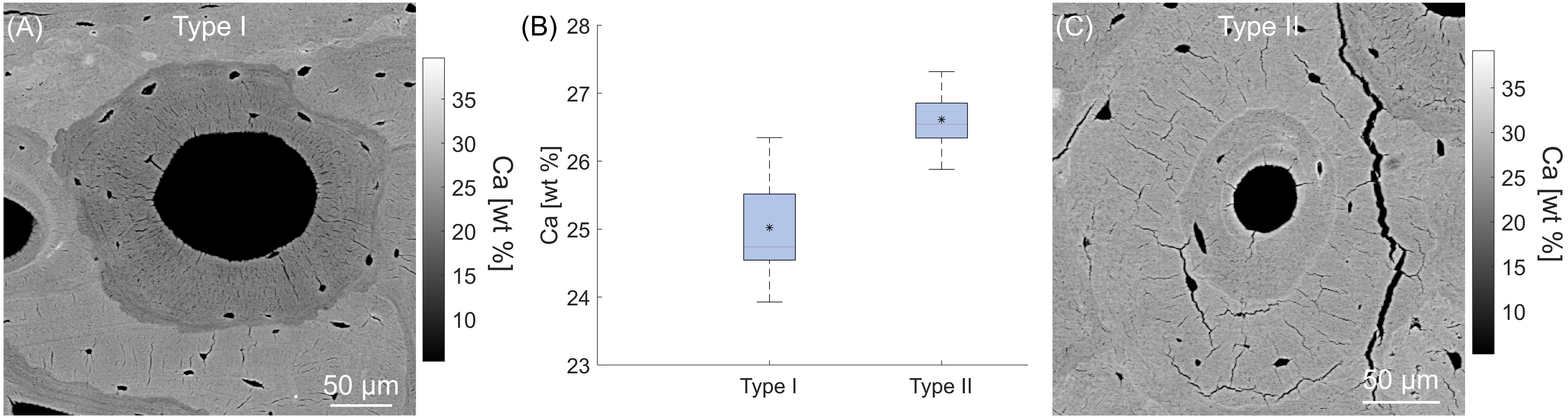

Fig. S6. Example of two osteons of different type and associated heterogeneity of the mineral content in the outer environment around the osteons. qBEI maps of the mineral content for (A) type I and (C) type II osteon (referred as osteon-in-osteon scenario). (B) Boxplot of the mean calcium content of the outer layer O2 around the two osteons. The boxes represent the interquartile range, encompassing 50% of the data, with the lower quartile at the bottom and the upper quartile at the top. The median and mean values are indicated by a line and an asterisk, respectively. The whiskers extend to show the range of data, reaching from the 5th percentile to the 95th percentile.

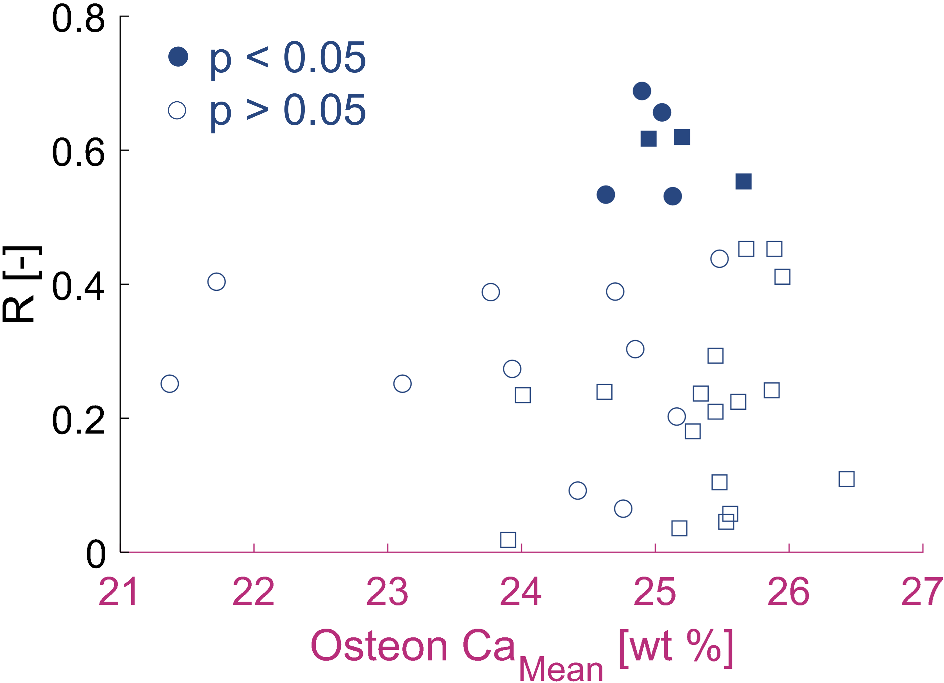

Fig. S7. Spearman correlation coefficients between the mineral content of the CL and inner layer I1, computed according to a spatially resolved analysis around each osteon and represented according to the statistical significance of their correlation. Circles represent osteons from the adult individual (40 years) and square from the aged individual (81 years).

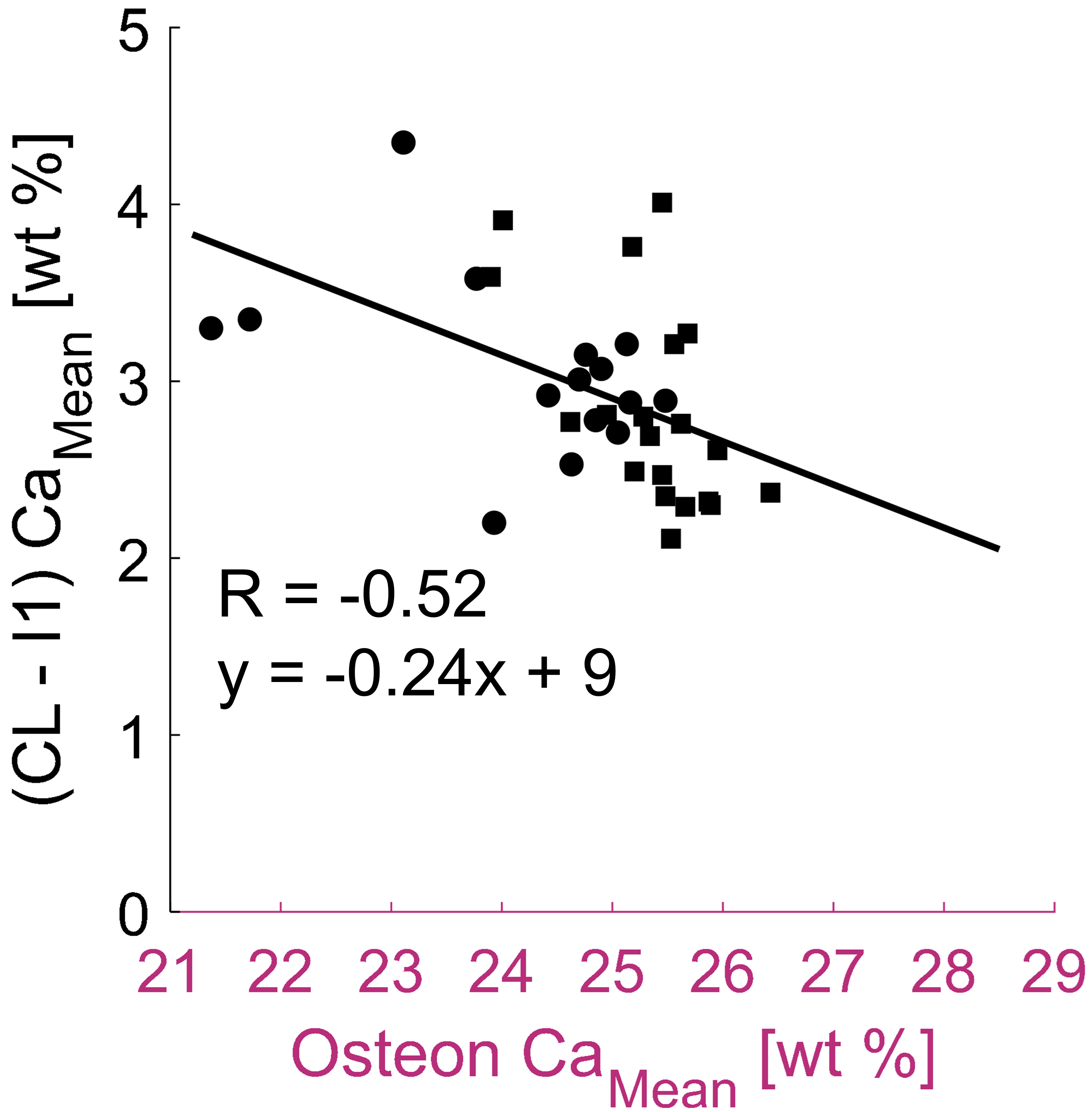

Fig. S8. Difference in mineral content between CL and closest neighboring environment inside the osteon (represented by the first inner layer I1), computed as average difference (CL-I1) in each osteon and plotted versus the mean degree of mineralization of the osteon. Circles represent osteons from the adult individual (40 years) and square from the aged individual (81 years).

To understand the interplay between primary and secondary mineralization, it is instructive to introduce a mathematical description of the mineralization law, quantifying the increase in calcium content *Ca* as a function of time *t* (elapsed from osteoid deposition) expressed in units of turnover time (which is defined as the time it takes to remodel an amount of bone equal to the actual bone volume). In trabecular bone, the mineralization law can be well-represented as a sum of two hyperbolic growth functions [1]:

$Ca \left( t \right)=C_{1}\frac{t/t_{1}}{1+t/t_{1}}+C_{2}\frac{t/t_{2}}{1+t/t_{2}}$	(S1)

with $C_{1}$, $t_{1}$ and $C_{2}$, $t_{2}$ being parameters describing primary and secondary mineralization, respectively. We assume that such analytical description is also valid for osteonal bone, but most likely with different parameters [2] and with a different turnover time [3]. Starting from a reference mineralization law, we explore the impact of an increased primary mineralization, with both $C_{1}$ and $t_{1}$ varied by 5, 10, 15 and 20% (Fig. S9A). Without modifying the secondary mineralization, the final mineral content attained by CL is higher than experimentally measured. Such behavior is not seen when, following a higher primary mineralization, there is a slower secondary mineralization, as detected in this study (Fig. S9B).

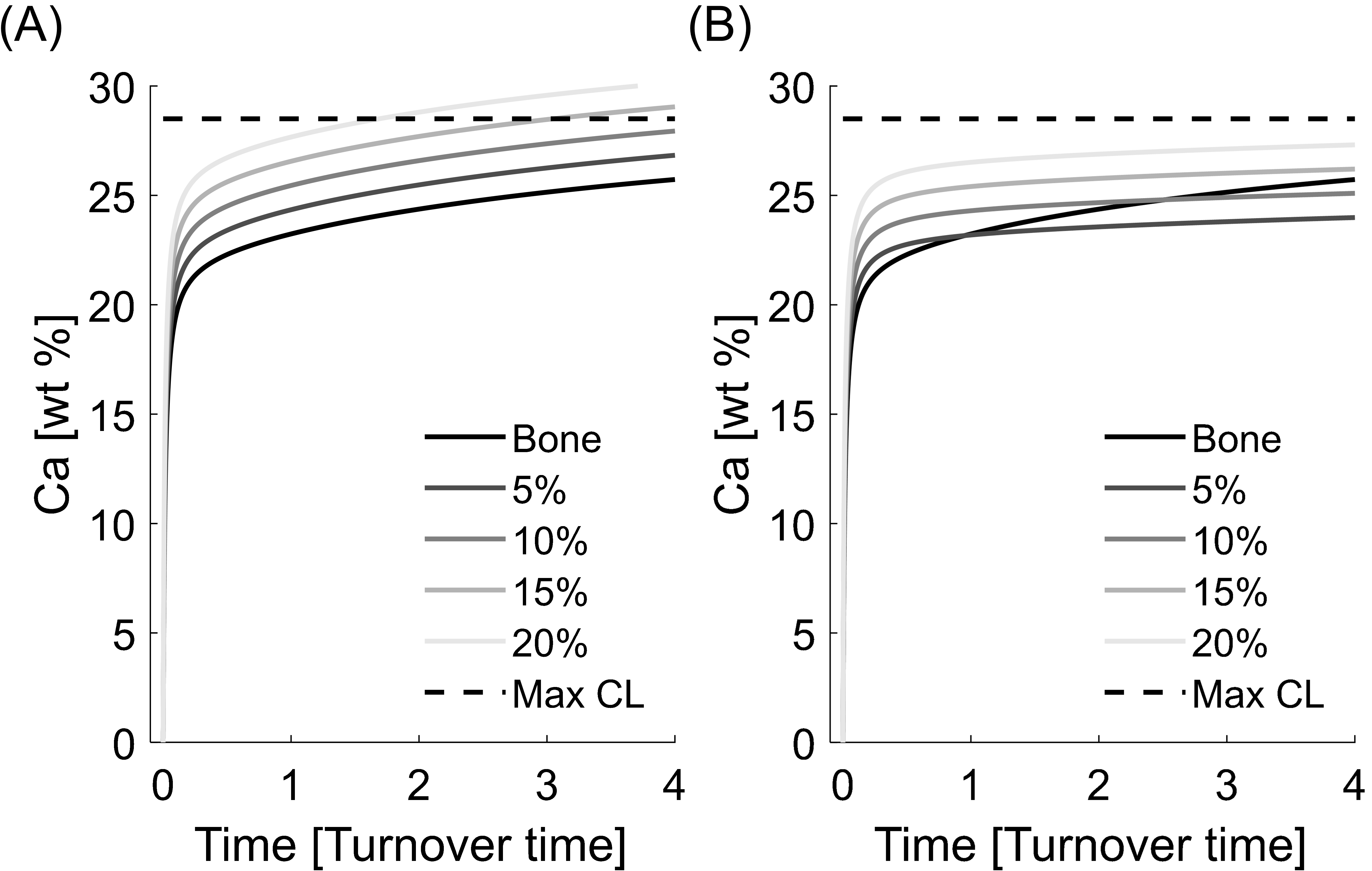

Fig. S9. Mathematical description of the mineralization process. (A) Mineralization laws of CL assuming an increased primary mineralization by 5%, 10%, 15% and 20%. (B) Mineralization laws of CL assuming an increased primary mineralization (by 5%, 10%, 15% and 20%) and a decreased secondary mineralization (by a factor of 2.5) with respect to the mineralization of the bone. The dotted line is the maximum mineral content measured in the CL in this work.

References

[1] Ruffoni D, Fratzl P, Roschger P, Klaushofer K, Weinkamer R. The bone mineralization density distribution as a fingerprint of the mineralization process. Bone. 2007 May;40(5):1308–19

[2] Lerebours C, Weinkamer R, Roschger A, Buenzli PR. Mineral density differences between femoral cortical bone and trabecular bone are not explained by turnover rate alone. Bone Rep. 2020 Dec;13:100731.

[3] Ericksen M. Histologic estimation of age at death using the anterior cortex of the femur. J. Phys. Anthropol. 1991 Dec;84(2):171–9.
